# ROCK2 Knockout Improves Proliferation Rate in a Cellular Model of Down Syndrome

**DOI:** 10.1101/2022.10.20.513071

**Authors:** Janine LeBlanc-Straceski, Riley Williams, Kylie Ward, Autumn Bates, Catherine Duran, Lordina Anderson, Morgan Murray, Jennifer Pereira-Badji, Christina Shoushani, Jacob Thibault

## Abstract

In a cellular model of Down Syndrome, hTERT immortalized RPE-1 (human retinal pigment epithelial-1) cells carrying an extra copy of chromosome 21 exhibit reduced fitness, in part, as an increase in doubling time (or a reduction in cell proliferation rate) in culture. ROCK2 (Rho associated coiled-coil containing kinase 2) was identified in a whole genome CRISPR knockout (KO) screen designed to identify genes and pathways that could be therapeutically targeted to improve cell proliferation (Replogle JM, et al. m*anuscript in preparation*). ROCK2 KO cell lines of both RPE-1 euploid and trisomy 21 aneuploid cells were created using CRISPR. Trisomy 21 ROCK2 KO cell lines showed a modest increase in cell proliferation rate compared to the parental aneuploid cells, similar to the relative effect that ROCK2 knockout had in the whole genome CRISPR screen. Euploid ROCK2 KO cell lines showed no difference in growth rate vs their ROCK2 expressing counterparts. Changes in doubling time in response to two pharmaceutical ROCK inhibitors, Fasudil and Y27632, also showed the same modest increase in cell proliferation rate in the trisomy 21 cells. The actin cytoskeleton, a target of ROCK2 regulation, exhibited long stress fibers that aligned across multiple contiguous cells in confluent trisomy 21 ROCK2 KO cells compared to the disorganized stress fibers of the parental trisomy 21 cells with normal ROCK2 expression.

## INTRODUCTION

Many different altered phenotypes are associated with human trisomy 21 or Down Syndrome (Antonarakis SE, et al. 2020) including hypocellularity (Guidi S, et al. 2011). Analysis of aneuploid cells carrying a third copy of chromosome 21 reveal that a reduction in cell proliferation rate versus euploid counterparts is one measure of decreased cell fitness (Contestabile A, et al. 2009; Hwang S, et al. 2019; Hwang S et al. 2021; Inoue M, et al. 2019; Gimeno A, et al. 2014; Guidi S, et al. 2011; Meharena HS, et al. 2022; Soppa U, et al. 2014; Stingele S, et al. 2012; Williams BR, et al. 2008). A whole genome CRISPR screen was undertaken in order to identify genes and pathways that would improve trisomy 21 cell proliferation and that could be targets for therapeutic intervention (Replogle JM, et al. m*anuscript in preparation*). One gene knockout that was highly enriched in the screen was Rho associated coiled-coil containing kinase 2 (ROCK2).

Two Rho associated kinase genes are found in the human genome: ROCK1 and ROCK2 on chromosomes 18 and 2, respectively. Both kinases were identified in the screen, but ROCK1 had a much lower enrichment score. Many studies on the different roles of these two regulatory kinases have produced a large body of literature describing many different cellular roles (Julian L, Olson, MF. 2014). ROCK2 is a serine/threonine protein kinase which, by phosphorylating myosin light chain 2 (Kawano Y, et al. 1999), remodels the actin cytoskeleton, including stress fiber and focal adhesion formation, and effects cell polarity (Amano M, et al. 2010). It also regulates vascular smooth muscle cell contraction (Hartmann S, et al. 2015; Kassianidou E, et al. 2017; Li Y, et al. 2021). In neurons, ROCK1 forms stable actomyosin filament bundles that initiate polarity in cells and dendritic spines. In contrast, ROCK2 regulates contractile force at the leading edge of migratory cells and regulates cofilin-mediated actin remodeling of adhesions (Newell-Litwa KA, et al., 2015). Several reports in the literature also mention the effects of ROCK function and MLCK in contractile ring formation and function, but fail to distinguish between the two human isoforms (Chircop M. 2017).

ROCK2 is a therapeutic target for many human disease conditions (Hartmann S, et al. 2015; Sharma P, Roy K. 2020; Weber AJ, Herskowitz JH. 2021), for example, the fibrosis associated with diabetic kidney disease (Nagai Y, et al. 2019) and graft versus host disease (cGVHD). A few key ROCK inhibitors, including Fasudil and Y27632 have been used extensively to study the roles of these two kinases. Recently, the FDA has approved a new ROCK2 specific inhibitor, belumosudil (REZUROCK, Kardamon, Sanofi) for the treatment of the fibrosis associated with cGVHD that occurs in some bone marrow transplant patients (Zanin-Zhorov A, Blazar BR. 2021). This action of ROCK2 is a completely different mechanism involving ROCK2 interaction with STAT3 in the cytosol, the formation of a ROCK2/JAK2/STAT3 complex, STAT3 phosphorylation and translocation to the nucleus where it regulates gene expression in T-cells of the immune system (Chen W, et al. 2018). This report highlights the need to assess ROCK function in a cell and system specific manner.

Few reports in the literature link ROCK2 activity to cell proliferation. Deletion of both ROCK1 and 2 blocked tumor formation and cell cycle progression (Kümper S, et al. 2016), and ROCK2 overexpression has been implicated in the increased proliferation of liver cancers (Zhang L, et al. 2019). Here we present evidence that in RPE-1 (retinal pigment epithelial-1) trisomy 21 cells, both ROCK2 CRISPR knockout and ROCK drug inhibition result in a modest but significant increase in the proliferation rate. CRISPR was used to knockout ROCK2 in hTERT immortalized aneuploid RPE-1 cells, which were created by microcell mediated chromosome transfer and generously shared by the Storchova lab (Stingele S, et al. 2012), and further modified by infection with a Cas9 expressing viral vector (Replogle JM, et al. m*anuscript in preparation*). Plasmids expressing the two most enriched sgRNAs for ROCK2 identified in the whole genome screen were transfected into the cells to knockout the gene. Comparison of the growth curves revealed that the trisomy 21 ROCK2 knockout cell lines had an increased cell proliferation rate over the aneuploid cells that retained the ROCK2 gene, while euploid cell lines were unaffected by knocking out ROCK2 expression. Corroborating results were achieved using Fasudil and Y27632, two ROCK inhibitors.

## MATERIALS AND METHODS

### Cell Lines

H-TERT immortalized RPE-1 H2B-GFP 21/3 cells were originally produced by microcell mediated chromosome transfer by the Storchova lab (Stingele, et al. 2012; Passerini, et al. 2016) and generously shared with us. This cell line and the parental H-TERT immortalized euploid RPE-1 cell line were infected to express Cas9 using LentiCas9-Blast (Sanjana NE, et al. 2014), and subsequently used in the whole genome CRISPR screen as described (Replogle JM, et al. m*anuscript in preparation*). Briefly, LentiCas9-Blast plasmid (a gift of Feng Zhang, Addgene plasmid # 52962; http://n2t.net/addgene:52962; RRID:Addgene_52962) was used to produce virus in 293T cells which was used to infect RPE-1 and RPE-1-21/3 cell lines. RPE-1-Cas9 and RPE-1 21/3-Cas9 cell lines (henceforth referred to as RPE1 WT-Cas9, and RPE1 21/3-Cas9) and the ROCK2 CRISPR knockout cell lines (described below) were grown in Dulbecco’s Modified Eagle Medium (DMEM; Invitrogen 11995) supplemented with 10% Fetal Bovine Serum, 1% Penicillin-Streptomycin, and 1% L-glutamine and passaged every three days.

### CRISPR sgRNA Vector Design

A VectorBuilder platform (https://en.vectorbuilder.com/design.html) was used to design plasmids to express the top two ROCK2 single guide RNAs (sgRNA-A: GTAGGTAAATCCGATGAA and sgRNA-B: GAGGTTCTGAAATCACAA) identified previously from the whole genome CRISPR knockout screen (Replogle JM, et al. m*anuscript in preparation*). These vectors also expressed an EGFP:P2A:Puro fusion protein for both puromycin selection and visual confirmation of a successful transfection by viewing cytoplasmic GFP expression (Fig. 1A, 1B).

**Fig. 1.**
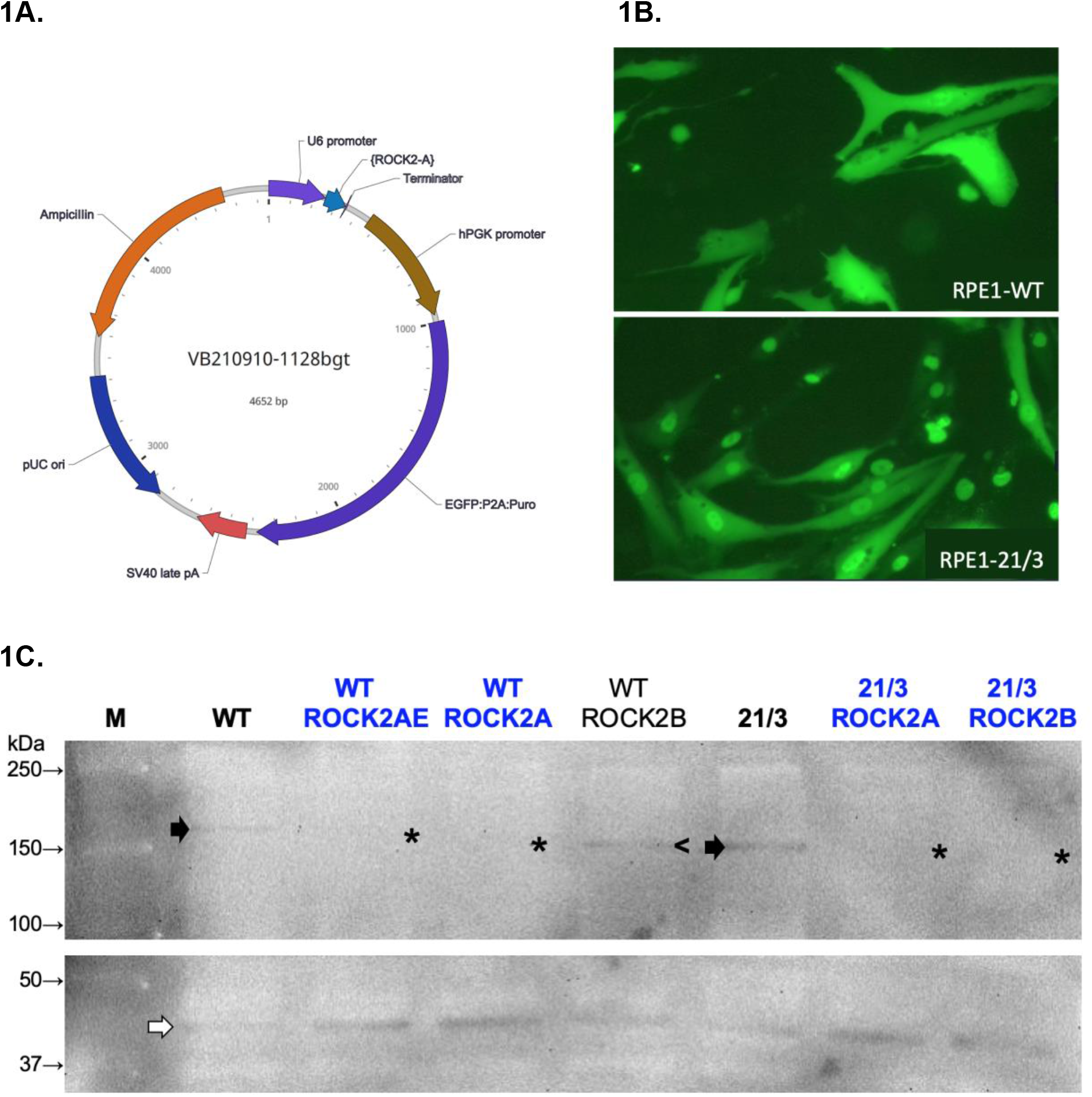
Creation of CRISPR RPE1 ROCK2 KO Cell lines. **A**. Vector map of the sgRNA Expression Vector highlighting the ROCK2A CRISPR sgRNA driven by a U6 promoter and the EGFP and Puromycin resistance coding regions linked by P2A driven by the hPGK promoter. A vector with the ROCK2B sgRNA was similarly generated. **B**. Cytoplasmic GFP expression of the EGFP:Puro fusion protein 24 hr post transfection of the RPE1 WT-Cas9 and RPE1 21/3-Cas9 cells. Note that the RPE1-21/3 cells (lower panel) carry a histone H3:GFP fusion and thus both the cytoplasm and nuclei of transfected cells appear green. **C**. Western Blot confirmation of CRISPR knockout of ROCK2. Antibody detection of ROCK2 shows that expression was eliminated in transfectants with ROCK2 sgRNA A in WT and 21/3 cells, and with ROCK2 sgRNA B in 21/3 cells (asterisks*) compared to untransfected controls (black arrows). Note that in one cell line tested labeled WT ROCK2B (lane 5), knockout of ROCK2 was unsuccessful (**<**). Actin loading controls are shown in bottom panel (white arrow).

### ROCK2 CRISPR Knockouts

The RPE1 WT-Cas9 and RPE1 21/3-Cas9 cells were transfected separately with either of the two ROCK2 sgRNA expressing plasmids using standard Invitrogen Lipofectamine 3000 protocol. After 24 hrs, a Zeiss Primovert inverted microscope was used to detect fluorescence from the EGFP:P2A:Puro fusion protein expression (Fig. 1B). Cells were selected in puromycin at 15 µg/ml for 7 days to eliminate non-transfected cells.

### Western Blot

Western blot analysis was used to confirm CRISPR knockout in cells transfected with ROCK2 sgRNA A or B expression vectors. RPE1 WT-Cas9 and RPE1 21/3-Cas9 transfected cells (3×10^6^ cells) were plated and collected on the following day. Cells were lysed in RIPA buffer (Thermo Scientific 89900) containing protease inhibitors, *cOmplete, Mini, EDTA-free Protease Inhibitor Cocktail (Roche)* and Sigma Fast Protease Inhibitor Tablets, S8820-20TAB. Protein concentration was determined using BioRad Bradford Protein Assay reagent. Protein was mixed with 2X Laemmli Buffer (Sigma-Aldrich 53401), run on BioRad 4-20% precast SDS-polyacrylamide gels, and transferred to Immobilon-P using a BioRad Semi-dry transfer apparatus. The blot was blocked in 3% milk in TBST, washed, then incubated in a 1:5000 dilution for both a ROCK2 Polyclonal Antibody (Invitrogen PA5-78291) and an anti-beta actin antibody (Proteintech 20536-1-AP). The blot was washed, re-blocked, and incubated with 1:2000 dilution of Invitrogen Goat anti-Rabbit IgG (H+L) Secondary Antibody, HRP conjugate (#31460). Chemiluminescence (Pierce ECL substrate) was detected using a Syngene G-box.

### Growth Curves

RPE1 WT-Cas9 or RPE1 21/3-Cas9 cells and their ROCK2 knockout (KO) counterparts were plated in 5 ml media in each of 2, T25(25 cm^2^ surface) flasks at a density of 6×10^3^ cells/cm^2^ cells or 1.5×10^5^ cells per flask. Every 3 days for 12 days, the cells were trypsinized, collected, counted in duplicate on a Cellometer Mini (Nexcelom) and replated at 1.5×10^5^ cells/flask. Growth rates were calculated using the formula: rate = (ln(N_t_/N_0_))/t, where N_t_ equals the number of cells at time = t, and N_0_ equals the number of cells plated at time t=0, and plotted using Google Sheets.

### ROCK Inhibitors

Cells were exposed to two different Rho Kinase (ROCK) inhibitors, Fasudil-HCl (Sigma-Aldrich) and Y27632 (EMD Millipore). RPE1 WT-Cas9 and RPE1 21/3-Cas9 cell lines were exposed to either of the two ROCK inhibitors at a concentration of 10µM of Fasudil or 250nM of Y-27632 in DMEM media. Drug concentrations were determined in accordance with the product specifications and previous studies (So, S., et. al 2020). Fresh inhibitors were introduced in the media during passaging every three days and cells were maintained in media containing the drug inhibitors throughout the 12 day growth curve protocol.

### Confocal Imaging

Cells were grown on MatTek 1.5 coverglass bottom 35 mm petri dishes, fixed in 4% paraformaldehyde, and permeabilized with 0.1% TX100 in PBS. Cells were stained with ActinRed ReadyProbe (Invitrogen). ProLong™ Gold Antifade Mountant with DAPI Blue (ThermoFisher) was used to stain the nuclei and preserve cells. Confocal imaging was performed on a Zeiss LSM 800 microscope using ZEN software. Confluent cell images were collected using uniform settings for laser power and gain to allow comparison of different cell lines. All images were observed using the 20x objective.

## RESULTS

### CRISPR ROCK2 Knockout Cell Lines

RPE1 21/3-Cas9 cells have a decreased cell proliferation rate over their euploid counterparts (Replogle JM, et al. m*anuscript in preparation*). A whole genome CRISPR screen identified ROCK2 as a gene whose knockout was enriched at the end of a two-week cell culturing protocol to select for an increased rate of cell proliferation (Replogle JM, et al. m*anuscript in preparation*). To confirm the ROCK2 result, CRISPR was used to knockout the gene in the same RPE-1 Cas9 cells used in the screen, and cell proliferation rates were measured. Figure 1 shows the results of the successful creation of two RPE1 21/3-ROCK2 KO and one RPE1 WT-ROCK2 cell lines, using the top two guide ROCK2 sgRNAs identified in the screen - ROCK2A and ROCK2B. Figure 1A shows the map of the vector, which has an EGFP:Puro fusion to both select for transfectants and screen for the efficiency of transfection. Cytoplasmic EGFP expression was detection 24 hours after transfection (Fig. 1B), indicating a successful transfection. Note that the RPE1-21/3-Cas9 cells carry a histone H3:GFP fusion (Stingele S, et al. 2012) and thus both the cytoplasm and nuclei of transfected cells appear green in the aneuploid cells (Fig. 1B, lower panel). The knockout of ROCK2 was confirmed by Western blot analysis (Fig. 1C) in the following three cell lines: RPE1 21/3-ROCK2A, RPE1 21/3-ROCK2B, and RPE1 WT-ROCK2A. Note that in one cell line tested, RPE1 WT-ROCK2B, CRISPR knockout of ROCK2 was unsuccessful as the protein is detected (lane 5).

### ROCK2 Gene Knockouts in 21/3 Aneuploid Cells Exhibit Increased Cell Proliferation

To determine the effects of ROCK2 knockout on cell proliferation, growth curve experiments were performed. For each growth curve, cells were plated on day 0, incubated for three days, harvested, counted and replated at the same initial concentration. In two independent experiments (Fig. 2A and Fig. 2B), both trisomy 21 ROCK2 knockout cell lines showed a small but measurable and reproducible increase in cell proliferation rate (red and yellow lines) compared to the trisomy 21 ROCK2 expressing cells (blue lines). The 21/3-ROCK2B cell line had a slightly increased cell proliferation rate (20% decrease in doubling time, Fig. 2D and Table1), over 21/3-ROCK2A (16% decrease in doubling time) pointing to variability in the creation of the CRISPR KO cell lines. Notably, the WT-ROCK2A cell line showed a minor 1.4% decrease in cell proliferation rate (Fig. 2D, Table 1), indicating that knocking out this gene in euploid cells has a negligible effect on the rate of cell proliferation.

**Table 1.** Doubling Times for KO Cell Lines and ROCK Inhibitor Treated Cells. See Materials and Methods for growth conditions and sampling protocols.

|  | Average Doubling Time (hours) | % Decrease in Doubling Time (hours) |
| --- | --- | --- |
| **RPE1 KO Cell Lines** |  |  |
| 21/3-Cas9 | 39.9 |  |
| 21/3-ROCK2A | 34.6 | 13.4 |
| 21/3-ROCK2B | 33.2 | 20.0 |
| WT-Cas9 | 26.5 |  |
| WT-ROCK2A | 26.9 | -1.4 |
| **RPE1 Cas9 Cell Lines + ROCK Inhibitors** |  |  |
| 21/3-Cas9 | 37.2 |  |
| 21/3-Cas9, 10 μM Fasudil | 34.4 | 7.8 |
| 21/3-Cas9, 250 nM Y27632 | 31.9 | 14.2 |
| WT-Cas9 | 20.6 |  |
| WT-Cas9, 10 μM Fasudil | 25 | -21.3 |
| WT-Cas9, 250 nM Y27632 | 21.3 | -3.4 |

**Fig. 2.**
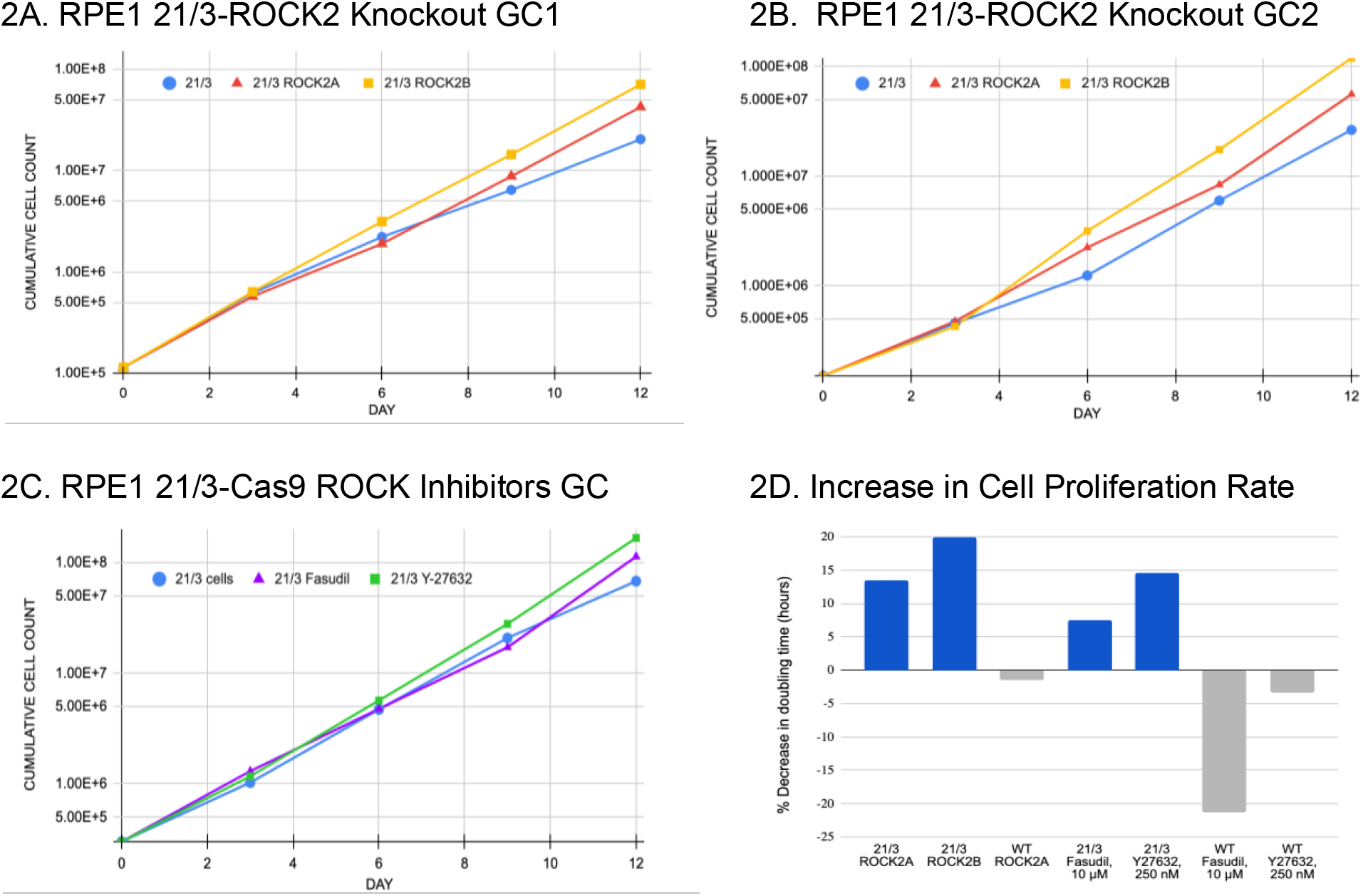
Cell Proliferation Rates in RPE1-21/3 Cells with ROCK2 Knockout and Drug Inhibitors. Cells were plated, grown for three days, then harvested, counted and replated for a total of 12 days (see Materials and Methods). **A+B**. Two separate growth curve experiments for the RPE1 21/3-Cas9 and CRISPR KO cell lines. Blue, 21/3-Cas9; Red, 21/3-ROCK2A, Yellow, 21/3-ROCK2B. **C**. Growth curves for the RPE1 21/3-Cas9 treated with ROCK inhibitors Fasudil (purple line, triangles) and Y27632 (green line, squares). **D**. The increase in cell proliferation rate, expressed as a percentage of decrease in doubling time relative to the control 21/3 or WT cell line, was calculated for ROCK2 knockout and inhibitor treated aneuploid and wild type cells. See Table 1.

### ROCK Inhibitors Increase Cell Proliferation Rate in Trisomy 21 Cells

Pharmaceutical inhibition of ROCK activity in aneuploid cell lines resulted in an increase in cell proliferation rate as indicated in Fig. 2C. The decrease in doubling time was comparable to the aneuploid ROCK2 knockouts as seen in Fig. 2A, 2B and Table 1. Cells treated with ROCK inhibitors displayed a similar slight increase in proliferation rate of 7.8% and 14.2 % for Fasudil and Y27632, respectively, similar to the approximately 17% average increase seen in the two aneuploid knockout cell lines (Table 1). This indicates that either knocking out or inhibiting ROCK2 in the RPE1 21/3-Cas9 cell lines leads to a small but consistent increase in the cell proliferation rate. In the WT-Cas9 cell line however, the drug inhibitors had opposite effects of variable magnitude. Y27632 had a negligible 3.4% increase in cell doubling time, very similar to the knockout WT-ROCK2A cell line, whereas Fasudil increased the doubling time by about 21% (Table 1).

### Actin Cytoskeleton

Myosin regulatory light chain is a target of the Rho dependent kinases, and ROCK2 in particular (Kawano Y, et al. 1999). To determine whether any cytoskeleton remodeling had taken place in the ROCK2 knockout cell lines, cells were grown to different densities and stained for actin. Figure 3 shows WT-Cas9 and 21/3-Cas9 plus the two aneuploid ROCK KO cell lines at low density. The actin cytoskeleton appears depleted in 21/3 cells compared to the WT counterparts. The aneuploid cells have less actin staining overall, and fewer stress fibers. Many of the aneuploid cells appear to have disorganized stress fibers, and the cells themselves are broad and flat. The ROCK2 KO cells appear have a slight increase in organized stress fibers, but the cells remain broad and flat.

**Fig 3.**
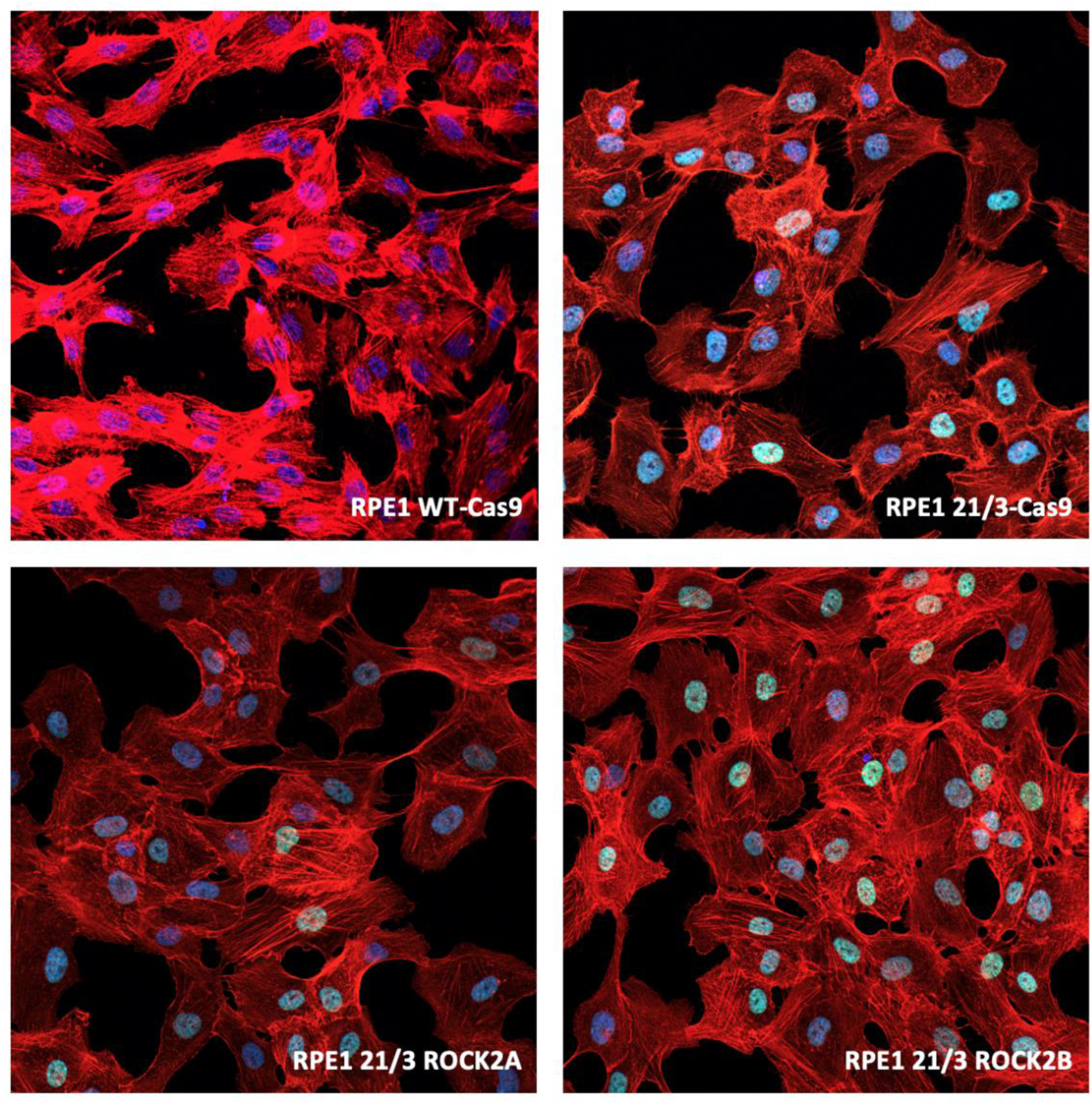
Actin Cytoskeleton in Low Density Cell Cultures. Cells were processed for imaging as described in the Materials and Methods. Note the RPE1 21/3 cells have a GFP-H3 fusion which renders the nuclei a teal color under dual DAPI and GFP fluorescence. 20x.

However, in KO cells that are grown to confluence, the number and organization of stress fibers is markedly different from the untreated counterparts (Fig. 4A). Figure 4 shows that both the 21/3-ROCK2A and -B cells appear to have a significant increase in number of organized stress fibers, which can be seen to align across multiple contiguous cells. (Fig. 4B).

**Fig 4.**
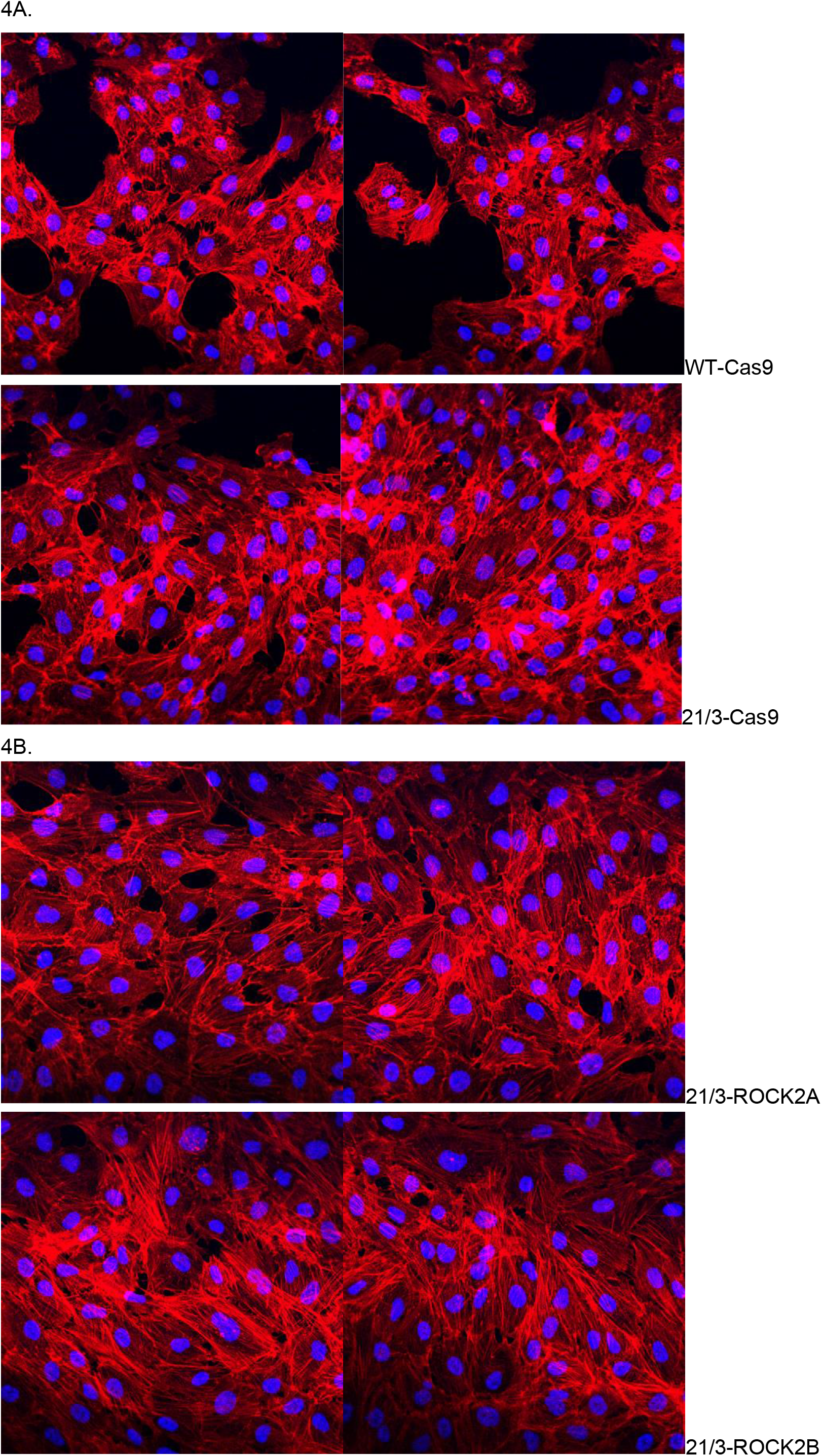
Actin Cytoskeleton in Confluent Cells. Cells were grown to confluence, processed for imaging as described in the Materials and Methods. **A**. Cells exhibiting typical stress fibers were seen in WT-Cas9 (top 2 panels) and 21/3-Cas9 cells (bottom 2 panels). **B**. Cells exhibiting long stress fibers that aligned across multiple contiguous cells were seen in 21/3 ROCK2A (top 2 panels) and 21/3 ROCK2B (bottom 2 panels).

## DISCUSSION

### ROCK2 Knockouts Confirm CRISPR Screen Results

Eliminating ROCK2 from trisomy 21 RPE1 cells caused a modest increase in their growth rate, comparable to the enrichment seen in the whole genome CRISPR screen (Replogle JM, et al. *manuscript in preparation)*. Therefore, these results support the overall design and execution of the CRISPR screen. They also present a challenge to provide an explanation for why knocking out this kinase increases the cell proliferation rate in the trisomy 21 cells. Since no increase was seen in the euploid cell ROCK2 knockouts, this is clearly a phenomenon related to the trisomy 21 genomic background. Although few reports have examined the role of ROCK knockouts in cell proliferation, contradictory reports exist of the effects on growth using ROCK inhibitors. ROCK inhibition has been shown to increase cell growth and recovery of cryopreserved stem cells (Claassen DA, et al. 2009). However, another report also claims that ROCK function is essential for cell cycle progression and tumorigenesis (Kümper S, et al. 2016). The involvement of ROCKs specifically, and kinases in general, in many aspects of cellular function requires careful study in many cell types. Nonetheless, the results presented here do demonstrate the validity of the CRISPR screen results.

### ROCK Inhibitors

The ROCK inhibitors selected for use in this study are commonly used in research and are prescribed to treat various ailments. When selecting the ROCK inhibitors, it was important to use an inhibitor that specifically targets ROCK2. Fasudil is a pharmaceutical inhibitor that is selective to ROCK2. It is being used clinically to treat cerebral vasospasm and for recovery care after stroke in order to improve cognitive decline (Chan S-L, Cipolla MJ. 2017). Although Y-27632 is not ROCK2 specific, it is the most commonly used ROCK inhibitor. Y-27632 has been used as a deterrent for apoptosis in iPSCs and other cell lines (Claassen DA, et al., 2009).

Published protocols for Fasudil demonstrated an optimal concentration between 1µM and 100µM in cell culture (So S, et al. 2020). A 10µM concentration was determined empirically for the RPE1 cells, since higher concentrations lead to rapid cell senescence (data not shown). A concentration of 250nM was used for the Y-27632 based on the manufacturer’s specifications. Trisomy 21 aneuploid cell lines treated with these inhibitors demonstrated an increase in proliferation rate when compared to the untreated aneuploid cell lines (Fig. 2C, D; Table 1), the magnitude of which was comparable to that seen in the trisomy 21 RPE1 ROCK2 KO cell lines. WT RPE1 cells treated with pharmaceutical inhibitors of ROCK2 however exhibited a decrease in proliferation rate, which was also similar to the results of the ROCK2 KO euploid cells (Fig. 2D; Table 1). This data supports the conclusion that when ROCK2 function is reduced or eliminated in trisomy 21 cells proliferation rates will improve. Therapeutic use of ROCK2 inhibitors such as belumosudil (REZUROCK, Kardamon, Sanofi) and Fasudil are already in clinical use for conditions associated with fibrosis in graft versus host disease (Ali, F et al. 2022; Cutler C, et al. 2021), in vasodilation in pulmonary hypertension (Christou H, et al. 2022), and for cerebral vasospasm associated with aneurysmal subarachnoid hemorrhage (Siasios, I et al. 2013).

The ROCK2/JAK2/STAT3 pathway, shown regulates gene expression in the nucleus of cells of the immune system, is another possible mechanism by which the depletion of ROCK2 could ameliorate cell proliferation in trisomy 21 cells (Chen W, et al. 2018). ROCK2 specific inhibitors have been shown to affect the transcription of downstream targets of this pathway (Flynn, et al. 2016). Further investigation should reveal whether ROCK2 enters the nucleus and influences gene expression patterns differentially in the trisomy 21 cells.

### Actin Cytoskeleton

The increase in stress fiber organization in the confluent ROCK2 KO trisomy 21 cells (Fig. 4) indicates a complex interaction between the Rho associate coiled-coil kinases and the stresses imposed on cells due to aneuploidy. In a study where both ROCK1 and ROCK2 isoforms were knocked down individually, the actin cytoskeleton was enhanced by ROCK2 down-regulation (Yoneda A, et al. 2005.). Fibroblasts with clear, robust stress fibers were seen when ROCK2 was knocked down using siRNA, however the general ROCK inhibitor Y27632 did not produce a similar effect. Cell proliferation rates were decreased when either or both ROCK isoforms were depleted. Other studies on ROCK2-/- MEFs (mouse embryonic fibroblasts) present images showing abundant organized, parallel stress fibers in cells in which ROCK2 was knocked down (Shi J, et al. 2013, Fig 4a), although the authors do not comment on this aspect of the figure in their report. Their conclusions are that ROCK1 can destabilize actin cytoskeleton by regulating MLC2 phosphorylation and thus peripheral actomyosin contraction, and that ROCK2 stabilizes actin cytoskeleton through regulating cofilin phosphorylation. Kümper S, et al. (2016) in Fig. 2A of their paper showed that cells with prominent stress fibers are seen in both the ROCK1 Δ/Δ and ROCK2 Δ/Δ cells. Cell proliferation rates moderately decreased when either ROCK isoform was knocked out, so there is not a correlation with the presence of fibers and increased cell proliferation rate. Cells could no longer perform cytokinesis when both ROCKs were knocked out simultaneously.

Clearly, aneuploid cells in general, and trisomy 21 cells in particular experience a different set of intracellular and environmental stresses than their euploid counterparts (Ben-David, U and Amon A, 2020). For example, aneuploidy has a profound influence on how cancer cells can mount different responses to chemotherapeutic agents (Replogle JM, et al. 2020). Aneuploid stress alters the intra- and extracellular environment of cells, and can influence the outcomes of the knockout CRISPR Screens which necessitates very careful interpretation of the results. However, it is also possible that this phenotype is specific to the trisomy 21 karyotype as opposed to other aneuploidies, driven by specific genes on chromosome 21. Pellegrini-Calace and Tramontano (2006) used bioinformatics to identify four chromosome 21 genes as putative members of a novel mitogen-activated protein kinase (MAPK) pathway. Of these genes - DYRK1A, SNF1LK, RIP4 and VPS26 endosomal protein sorting factor C (aka DSCR3 p) - only the first has been the subject of extensive investigation, though much more is needed. It is interesting to speculate that knocking out ROCK2 may mitigate the effects of the over expression of other kinases whose genes reside on chromosome 21, such as DYRK1a, which is another regulator of the actin cytoskeleton (Park J et al. 2012; Atas-Ozcan H, et al. 2021). Although there is no evidence that ROCK2 and DYRK1a interact directly or indirectly through a common pathway, they both ultimately regulate the actin cytoskeleton. Depleting ROCK2 may attenuate the effect of having three copies of DYRK1a, resulting in improved cell fitness. Ultimately, more work is needed to determine how ROCK2 knockout drives an increase in trisomy 21 proliferation and whether this observed effect of ROCK2 inhibition is trisomy 21-specific or broadly true for other aneuploid cells.

## ACKNOWLEDGEMENTS

Thanks to the Department of Biology in the School of Science and Engineering at Merrimack College for support of this work, Violet Tran for technical assistance, and Dr. John Replogle for a critical review of this manuscript. This project was initiated by Dr. John Replogle in the laboratory of Dr. Angelika Amon at MIT, whose presence will forever remain in our hearts, whose example remains a beacon to light our path forward, and whose contributions and human legacy will continue to enrich the collective body of scientific knowledge.

## REFERENCES

Ali F, Ilyas A. Belumosudil with ROCK-2 inhibition: chemical and therapeutic development to FDA approval for the treatment of chronic graft-versus-host disease. Curr Res Transl Med. 2022 Jul;70(3):103343. doi: 10.1016/j.retram.2022.103343. Epub 2022 Mar 24. PMID: 35339032.

Amano M, Nakayama M, Kaibuchi K. Rho-kinase/ROCK: A key regulator of the cytoskeleton and cell polarity. Cytoskeleton (Hoboken). 2010 Sep;67(9):545–54. doi: 10.1002/cm.20472. PMID: 20803696; PMCID: PMC3038199.

Antonarakis SE, Skotko BG, Rafii MS, Strydom A, Pape SE, Bianchi DW, Sherman SL, Reeves RH. Down syndrome. Nat Rev Dis Primers. 2020 Feb 6;6(1):9. doi: 10.1038/s41572-019-0143-7. PMID: 32029743; PMCID: PMC8428796.

Atas-Ozcan H, Brault V, Duchon A, Herault Y. Dyrk1a from Gene Function in Development and Physiology to Dosage Correction across Life Span in Down Syndrome. Genes (Basel). 2021 Nov 20;12(11):1833. doi: 10.3390/genes12111833. PMID: 34828439; PMCID: PMC8624927.

Ben-David U, Amon A. Context is everything: aneuploidy in cancer. Nat Rev Genet. 2020 Jan;21(1):44–62. doi: 10.1038/s41576-019-0171-x. Epub 2019 Sep 23. PMID: 31548659.

Chan SL, Cipolla MJ. Treatment with low dose fasudil for acute ischemic stroke in chronic hypertension. J Cereb Blood Flow Metab. 2017 Sep;37(9):3262–3270. doi: 10.1177/0271678x17718665. Epub 2017 Jun 30. PMID: 28665172; PMCID: PMC5584704.

Chen W, Nyuydzefe MS, Weiss JM, Zhang J, Waksal SD, Zanin-Zhorov A. ROCK2, but not ROCK1 interacts with phosphorylated STAT3 and co-occupies TH17/TFH gene promoters in TH17-activated human T cells. Sci Rep. 2018 Nov 9;8(1):16636. doi: 10.1038/s41598-018-35109-9. PMID: 30413785; PMCID: PMC6226480.

Chircop M. Rho GTPases as regulators of mitosis and cytokinesis in mammalian cells. Small GTPases. 2014;5:e29770. doi: 10.4161/sgtp.29770. Epub 2014 Jul 2. PMID: 24988197; PMCID: PMC4160341.

Christou H, Khalil RA. Mechanisms of pulmonary vascular dysfunction in pulmonary hypertension and implications for novel therapies. Am J Physiol Heart Circ Physiol. 2022 May 1;322(5):H702–H724. doi: 10.1152/ajpheart.00021.2022. Epub 2022 Feb 25. PMID: 35213243; PMCID: PMC8977136.

Claassen DA, Desler MM, Rizzino A. ROCK inhibition enhances the recovery and growth of cryopreserved human embryonic stem cells and human induced pluripotent stem cells. Mol Reprod Dev. 2009 Aug;76(8):722–32. doi: 10.1002/mrd.21021. PMID: 19235204; PMCID: PMC3257892.

Contestabile A, Fila T, Bartesaghi R, Ciani E. Cell cycle elongation impairs proliferation of cerebellar granule cell precursors in the Ts65Dn mouse, an animal model for Down syndrome. Brain Pathol. 2009 Apr;19(2):224–37. doi: 10.1111/j.1750-3639.2008.00168.x. Epub 2008 May 14. PMID: 18482164; PMCID: PMC8094641.

Cutler C, Lee SJ, Arai S, Rotta M, Zoghi B, Lazaryan A, Ramakrishnan A, DeFilipp Z, Salhotra A, Chai-Ho W, Mehta R, Wang T, Arora M, Pusic I, Saad A, Shah NN, Abhyankar S, Bachier C, Galvin J, Im A, Langston A, Liesveld J, Juckett M, Logan A, Schachter L, Alavi A, Howard D, Waksal HW, Ryan J, Eiznhamer D, Aggarwal SK, Ieyoub J, Schueller O, Green L, Yang Z, Krenz H, Jagasia M, Blazar BR, Pavletic S. Belumosudil for chronic graft-versus-host disease after 2 or more prior lines of therapy: the ROCKstar Study. Blood. 2021 Dec 2;138(22):2278–2289. doi: 10.1182/blood.2021012021. Erratum in: Blood. 2022 Mar 17;139(11):1772. PMID: 34265047; PMCID: PMC8641099.

Flynn R, Paz K, Du J, Reichenbach DK, Taylor PA, Panoskaltsis-Mortari A, Vulic A, Luznik L, MacDonald KK, Hill GR, Nyuydzefe MS, Weiss JM, Chen W, Trzeciak A, Serody JS, Aguilar EG, Murphy WJ, Maillard I, Munn D, Koreth J, Cutler CS, Antin JH, Ritz J, Waksal SD, Zanin-Zhorov A, Blazar BR. Targeted Rho-associated kinase 2 inhibition suppresses murine and human chronic GVHD through a Stat3-dependent mechanism. Blood. 2016 Apr 28;127(17):2144–54. doi: 10.1182/blood-2015-10-678706. Epub 2016 Mar 16. PMID: 26983850; PMCID: PMC4850869.

Gimeno A, García-Giménez JL, Audí L, Toran N, Andaluz P, Dasí F, Viña J, Pallardó FV. Decreased cell proliferation and higher oxidative stress in fibroblasts from Down Syndrome fetuses. Preliminary study. Biochim Biophys Acta. 2014 Jan;1842(1):116–25. doi: 10.1016/j.bbadis.2013.10.014. Epub 2013 Oct 31. PMID: 24184606.

Guidi S, Ciani E, Bonasoni P, Santini D, Bartesaghi R. Widespread proliferation impairment and hypocellularity in the cerebellum of fetuses with down syndrome. Brain Pathol. 2011 Jul;21(4):361–73. doi: 10.1111/j.1750-3639.2010.00459.x. Epub 2010 Dec 6. PMID: 21040072; PMCID: PMC8094247.

Hartmann S, Ridley AJ, Lutz S. The Function of Rho-Associated Kinases ROCK1 and ROCK2 in the Pathogenesis of Cardiovascular Disease. Front Pharmacol. 2015 Nov 20;6:276. doi: 10.3389/fphar.2015.00276. PMID: 26635606; PMCID: PMC4653301.

Hwang S, Cavaliere P, Li R, Zhu LJ, Dephoure N, Torres EM. Consequences of aneuploidy in human fibroblasts with trisomy 21. Proc Natl Acad Sci U S A. 2021 Feb 9;118(6):e2014723118. doi: 10.1073/pnas.2014723118. PMID: 33526671; PMCID: PMC8017964.

Hwang S, Williams JF, Kneissig M, Lioudyno M, Rivera I, Helguera P, Busciglio J, Storchova Z, King MC, Torres EM. Suppressing Aneuploidy-Associated Phenotypes Improves the Fitness of Trisomy 21 Cells. Cell Rep. 2019 Nov 19;29(8):2473-2488.e5. doi: 10.1016/j.celrep.2019.10.059. PMID: 31747614; PMCID: PMC6886690.

Inoue M, Kajiwara K, Yamaguchi A, Kiyono T, Samura O, Akutsu H, Sago H, Okamoto A, Umezawa A. Autonomous trisomic rescue of Down syndrome cells. Lab Invest. 2019 Jun;99(6):885–897. doi: 10.1038/s41374-019-0230-0. Epub 2019 Feb 13. PMID: 30760866; PMCID: PMC6760570.

Julian L, Olson MF. Rho-associated coiled-coil containing kinases (ROCK): structure, regulation, and functions. Small GTPases. 2014;5:e29846. doi: 10.4161/sgtp.29846. Epub 2014 Jul 10. PMID: 25010901; PMCID: PMC4114931.

Kassianidou E, Hughes JH, Kumar S. Activation of ROCK and MLCK tunes regional stress fiber formation and mechanics via preferential myosin light chain phosphorylation. Mol Biol Cell. 2017 Dec 15;28(26):3832–3843. doi: 10.1091/mbc.E17-06-0401. Epub 2017 Oct 18. PMID: 29046396; PMCID: PMC5739298.

Kawano Y, Fukata Y, Oshiro N, Amano M, Nakamura T, Ito M, Matsumura F, Inagaki M, Kaibuchi K. Phosphorylation of myosin-binding subunit (MBS) of myosin phosphatase by Rho-kinase in vivo. J Cell Biol. 1999 Nov 29;147(5):1023–38. doi: 10.1083/jcb.147.5.1023. PMID: 10579722; PMCID: PMC2169354.

Kümper S, Mardakheh FK, McCarthy A, Yeo M, Stamp GW, Paul A, Worboys J, Sadok A, Jørgensen C, Guichard S, Marshall CJ. Rho-associated kinase (ROCK) function is essential for cell cycle progression, senescence and tumorigenesis. Elife. 2016 Jan 14;5:e12994. doi: 10.7554/eLife.12203. PMID: 26765561; PMCID: PMC4798951.

Li Y, Tai HC, Sladojevic N, Kim HH, Liao JK. Vascular Stiffening Mediated by Rho-Associated Coiled-Coil Containing Kinase Isoforms. J Am Heart Assoc. 2021 Oct 19;10(20):e022568. doi: 10.1161/JAHA.121.022568. Epub 2021 Oct 6. PMID: 34612053; PMCID: PMC8751888.

Meharena HS, Marco A, Dileep V, Lockshin ER, Akatsu GY, Mullahoo J, Watson LA, Ko T, Guerin LN, Abdurrob F, Rengarajan S, Papanastasiou M, Jaffe JD, Tsai LH. Down-syndrome-induced senescence disrupts the nuclear architecture of neural progenitors. Cell Stem Cell. 2022 Jan 6;29(1):116-130.e7. doi: 10.1016/j.stem.2021.12.002. PMID: 34995493; PMCID: PMC8805993.

Nagai Y, Matoba K, Kawanami D, Takeda Y, Akamine T, Ishizawa S, Kanazawa Y, Yokota T, Utsunomiya K, Nishimura R. ROCK2 regulates TGF-β-induced expression of CTGF and profibrotic genes via NF-κB and cytoskeleton dynamics in mesangial cells. Am J Physiol Renal Physiol. 2019 Oct 1;317(4):F839–F851. doi: 10.1152/ajprenal.00596.2018. Epub 2019 Jul 31. PMID: 31364374.

Newell-Litwa KA, Badoual M, Asmussen H, Patel H, Whitmore L, Horwitz AR. ROCK1 and 2 differentially regulate actomyosin organization to drive cell and synaptic polarity. J Cell Biol. 2015 Jul 20;210(2):225–42. doi: 10.1083/jcb.201504046. Epub 2015 Jul 13. PMID: 26169356; PMCID: PMC4508895.

Park J, Sung JY, Park J, Song WJ, Chang S, Chung KC. Dyrk1A negatively regulates the actin cytoskeleton through threonine phosphorylation of N-WASP. J Cell Sci. 2012 Jan 1;125(Pt 1):67–80. doi: 10.1242/jcs.086124. Epub 2012 Jan 16. PMID: 22250195.

Passerini V, Ozeri-Galai E, de Pagter MS, Donnelly N, Schmalbrock S, Kloosterman WP, Kerem B, Storchová Z. The presence of extra chromosomes leads to genomic instability. Nat Commun. 2016 Feb 15;7:10754. doi: 10.1038/ncomms10754. PMID: 26876972; PMCID: PMC4756715.

Pellegrini-Calace M, Tramontano A. Identification of a novel putative mitogen-activated kinase cascade on human chromosome 21 by computational approaches. Bioinformatics. 2006 Apr 1;22(7):775–8. doi: 10.1093/bioinformatics/btl006. Epub 2006 Jan 20. PMID: 16428264.

Replogle JM, LeBlanc-Straceski JM, Amon A. A Whole Genome CRISPR Screen to Relieve the Proliferation Defect of Trisomy 21 Cells. Manuscript in preparation.

Replogle JM, Zhou W, Amaro AE, McFarland JM, Villalobos-Ortiz M, Ryan J, Letai A, Yilmaz O, Sheltzer J, Lippard SJ, Ben-David U, Amon A. Aneuploidy increases resistance to chemotherapeutics by antagonizing cell division. Proc Natl Acad Sci U S A. 2020 Dec 1;117(48):30566–30576. doi: 10.1073/pnas.2009506117. Epub 2020 Nov 17. PMID: 33203674; PMCID: PMC7720170.

Sanjana NE, Shalem O, Zhang F. Improved vectors and genome-wide libraries for CRISPR screening. Nat Methods. 2014 Aug;11(8):783–784. doi: 10.1038/nmeth.3047. PMID: 25075903; PMCID: PMC4486245.

Sharma P, Roy K. ROCK-2-selective targeting and its therapeutic outcomes. Drug Discov Today. 2020 Feb;25(2):446–455. doi: 10.1016/j.drudis.2019.11.017. Epub 2019 Dec 12. PMID: 31837997.

Shi J, Wu X, Surma M, Vemula S, Zhang L, Yang Y, Kapur R, Wei L. Distinct roles for ROCK1 and ROCK2 in the regulation of cell detachment. Cell Death Dis. 2013 Feb 7;4(2):e483. doi: 10.1038/cddis.2013.10. PMID: 23392171; PMCID: PMC3734810.

Siasios I, Kapsalaki EZ, Fountas KN. Cerebral vasospasm pharmacological treatment: an update. Neurol Res Int. 2013;2013:571328. doi: 10.1155/2013/571328. Epub 2013 Jan 31. PMID: 23431440; PMCID: PMC3572649.

So S, Lee Y, Choi J, Kang S, Lee J-Y, Hwang J, Shin J, Dutton JR, Seo E-J, Lee BH, et al. 2020. The Rho-associated kinase inhibitor fasudil can replace Y-27632 for use in human pluripotent stem cell research. Bao X, editor. PLOS ONE. 15(5):e0233057. doi:10.1371/journal.pone.0233057.

Soppa U, Schumacher J, Florencio Ortiz V, Pasqualon T, Tejedor FJ, Becker W. The Down syndrome-related protein kinase DYRK1A phosphorylates p27(Kip1) and Cyclin D1 and induces cell cycle exit and neuronal differentiation. Cell Cycle. 2014;13(13):2084–100. doi: 10.4161/cc.29104. Epub 2014 May 7. PMID: 24806449; PMCID: PMC4111700.

Stingele S, Stoehr G, Peplowska K, Cox J, Mann M, Storchova Z. Global analysis of genome, transcriptome and proteome reveals the response to aneuploidy in human cells. Mol Syst Biol. 2012;8:608. doi: 10.1038/msb.2012.40. PMID: 22968442; PMCID: PMC3472693.

Weber AJ, Herskowitz JH. Perspectives on ROCK2 as a Therapeutic Target for Alzheimer’s Disease. Front Cell Neurosci. 2021 Mar 15;15:636017. doi: 10.3389/fncel.2021.636017. PMID: 33790742; PMCID: PMC8005730.

Williams BR, Prabhu VR, Hunter KE, Glazier CM, Whittaker CA, Housman DE, Amon A. Aneuploidy affects proliferation and spontaneous immortalization in mammalian cells. Science. 2008 Oct 31;322(5902):703–9. doi: 10.1126/science.1160058. PMID: 18974345; PMCID: PMC2701511.

Yoneda A, Multhaupt HA, Couchman JR. The Rho kinases I and II regulate different aspects of myosin II activity. J Cell Biol. 2005 Aug 1;170(3):443–53. doi: 10.1083/jcb.200412043. Epub 2005 Jul 25. PMID: 16043513; PMCID: PMC2171463.

Zanin-Zhorov A, Blazar BR. ROCK2, a critical regulator of immune modulation and fibrosis has emerged as a therapeutic target in chronic graft-versus-host disease. Clin Immunol. 2021 Sep;230:108823. doi: 10.1016/j.clim.2021.108823. Epub 2021 Aug 14. PMID: 34400321; PMCID: PMC8456981.

Zhang L, Zhou H, Wei G. miR-506 regulates cell proliferation and apoptosis by affecting RhoA/ROCK signaling pathway in hepatocellular carcinoma cells. Int J Clin Exp Pathol. 2019 Apr 1;12(4):1163-1173. PMID: 31933931; PMCID: PMC6947048.

